# Benchmarking GE-BOLD, SE-BOLD, and SS-SI-VASO sequences for depth-dependent separation of feedforward and feedback signals in high-field MRI

**DOI:** 10.1101/2021.12.10.472064

**Authors:** Polina Iamshchinina, Daniel Haenelt, Robert Trampel, Nikolaus Weiskopf, Daniel Kaiser, Radoslaw M. Cichy

## Abstract

Recent advances in high-field fMRI have allowed differentiating feedforward and feedback information in the grey matter of the human brain. For continued progress in this endeavor, it is critical to understand how MRI data acquisition parameters impact the read-out of information from laminar response profiles. Here, we benchmarked three different MR-sequences at 7T - gradient-echo (GE), spin-echo (SE) and vascular space occupancy imaging (VASO) - in differentiating feedforward and feedback signals in human early visual cortex (V1). The experiment (N=4) consisted of two complementary tasks: a perception task that predominantly evokes feedforward signals and a working memory task that relies on feedback signals. In the perception task, participants saw flickering oriented gratings while detecting orthogonal color-changes. In the working memory task, participants memorized the precise orientation of a grating. We used multivariate pattern analysis to read out the perceived (feedforward) and memorized (feedback) grating orientation from neural signals across cortical depth. Analyses across all the MR-sequences revealed perception signals predominantly in the middle cortical compartment of area V1 and working memory signals in the deep compartment. Despite an overall consistency across sequences, SE-EPI was the only sequence where both feedforward and feedback information were differently pronounced across cortical depth in a statistically robust way. We therefore suggest that in the context of a typical cognitive neuroscience experiment as the one benchmarked here, SE-EPI may provide a favorable trade-off between spatial specificity and signal sensitivity.

**Highlights:** Here, we benchmarked three sequences at high-field fMRI -GE-BOLD, SE-BOLD and VASO - in differentiating feedforward and feedback signals across grey matter depth of area V1. We show that:

- All the MR-sequences revealed the feedforward and feedback signals at the middle and deep cortical bins, respectively.
- Such correspondence across the sequences indicates that widely used GE-BOLD is a suitable method for the exploration of signals in cortical depth.
- Only SE-BOLD yielded statistically reliable differences between the cortical bins carrying feedforward and feedback signals.

## 1. Introduction

Recent advances in high-field fMRI have allowed neuroscientists to differentiate feedforward and feedback signals across cortical depth in the healthy human brain. Feedback signals were identified in superficial and/or deep grey matter bins, whereas feedforward signals were found at the middle cortical depth or across all the cortical bins (Bergmann et al., 2019; Lawrence et al., 2018; Iamshchinina et al., 2021). Such depth-specific separation of feedforward and feedback signals has been most firmly established in primary visual cortex (Muckli et al., 2015; Kok et al., 2016; Lawrence et al., 2018; Bergmann et al., 2019; Aitkens et al., 2021; Iamshchinina et al., 2021), but has also been found in somatosensory cortex (Yu et al., 2019). In most of these studies, the depth-specific signal differentiation was established using gradient-echo echo-planar imaging sequence (GE-BOLD). While GE-BOLD yields strong blood-oxygen level dependent (BOLD) responses and thus offers high signal sensitivity, it is argued to offer relatively low spatial specificity (Menon et al., 1995; Turner, 2002; Koopmans et al., 2010; Polimeni et al., 2010; Gagnon et al., 2016). As spatial specificity is essential for the precise estimation of depth-dependent cortical profiles, alternative acquisition methods that offer higher spatial specificity have recently attracted attention: particularly, spin-echo (SE)-EPI BOLD (Yacoub et al., 2003; Duong et al., 2003; Olman et al., 2010; Boyacioglu et al., 2014) and the cerebral blood volume (CBV)-based laminar fMRI using the SS-SI-vascular space occupancy (VASO) method (Huber et al., 2014). On the flipside, however, these methods yield reduced signal sensitivity compared to GE-BOLD (Zhao et. al., 2006; Huber et al., 2016a). As the problem of signal differentiation in grey matter depth requires both high spatial specificity and signal sensitivity, it is unclear how the tradeoff offered by each of the imaging methods pans out in a typical cognitive neuroscience experiment targeting feedforward and feedback information readout at 7T fMRI.

Here, we benchmarked three MR-sequences at 7T - GE-BOLD, SE-BOLD and SS-SI-VASO - in their ability to differentiate feedforward and feedback signals in human early visual cortex (V1). The experiment consisted of four participants completing multiple scanning sessions with two complementary tasks: a perception task that predominantly probed cortical feedforward signals, and a working memory task that engages feedback signals. In the perception task, participants viewed oriented gratings while reacting to color changes of fixation cross. In the working memory task, they had to maintain specific grating orientations in working memory. We used multivariate pattern analysis to read out the perceived (feedforward) and memorized (feedback) orientations from neuronal signals across cortical depth. Basing our hypotheses on previous layer-specific work, we expected the perception signals to be predominantly encoded in the middle cortical bin and the working memory signals to emerge in one or both outer cortical bins (superficial and deep). We estimated these effects in every MR-sequence. For comparability of the results across the three imaging methods, we performed equivalent post-processing steps on the acquired data and estimated the effects on closely matching time intervals in the trial.

Data analysis across all the MR-sequences revealed the feedforward signal predominantly at the middle cortical depth bin of area V1 and the feedback signal in the deep bin. The results obtained with conventional GE-BOLD generally agree with the results obtained with VASO and SE-BOLD, indicating that GE-BOLD is a suitable method for the exploration of signals in cortical depth with 7T fMRI. The VASO sequence showed weak representations of both feedforward and feedback signals, although the overall pattern of results corresponded to those found for the other sequences. Interestingly, SE-BOLD was the only sequence yielding statistically reliable differences between the cortical bins that carry feedforward and feedback signals. We therefore suggest that in the context of a typical cognitive experiment such as here, SE-BOLD may provide a suitable trade-off between spatial specificity and signal sensitivity.

## 2. Methods

### 2.1. Participants

Four adults (age in range from 28 to 35 years; 2 female) participated in the study. All participants had normal or corrected-to-normal vision. Each participant had extensive experience with participating in psychophysics studies at high-field fMRI and participated in two sessions per MR-sequence (GE, SE, VASO), resulting in 6 sessions per participant and 24 sessions in total. Participants gave their written informed consent for participation in the study as well as for publicly sharing all obtained data in pseudonymized form. They received monetary reimbursement for their participation. One of the participants did not complete all the perception runs during the first session of GE and first session of SE measurements, and thus only the data from the second sessions of both sequences is utilized for the feedforward signal estimation. The study was approved by the ethics committee of the Medical Faculty, University of Leipzig, Germany.

### 2.2. Stimuli

Stimuli were grayscale luminance-defined sinusoidal gratings generated using MATLAB (MathWorks, Natick, MA) with the Psychophysics Toolbox (Brainard, 1997). The gratings were presented in an annulus (outer diameter: 6.7° of visual angle, inner diameter: 1.3° of visual angle) surrounding a central fixation point. The gratings had a spatial frequency of 2 cpd and a Michelson contrast of 50%. Stimuli were displayed on an LCD projector (Sanyo PLC-XT20L) positioned in front of the scanner within the magnet room. Participants viewed the screen through a mirror attached to the head coil.

### 2.3. Experimental procedure

#### 2.3.1. Training procedure

Before entering the MRI scanner, participants underwent a training which comprised a minimum of 4 runs for all the participants. At the start of each trial, participants briefly saw two randomly oriented gratings (Figure 1). A subsequently presented task cue indicated which of the gratings needed to be memorized for the upcoming task. After an 11 second retention time a probe grating was shown. The grating was slightly tilted clockwise or counterclockwise with respect to the grating that had been cued; the amount of additional tilt was regulated in a staircase procedure (described below). The participants’ task was to indicate whether the probe grating was tilted clockwise or counterclockwise from the memorized grating. After each trial, participants received a 1 second feedback about their response correctness. The inter-trial interval was 2 seconds. Each training run consisted of 16 trials and took 4 minutes 54 seconds. At the end of each run, participants received feedback about their average accuracy.

**Figure 1.**
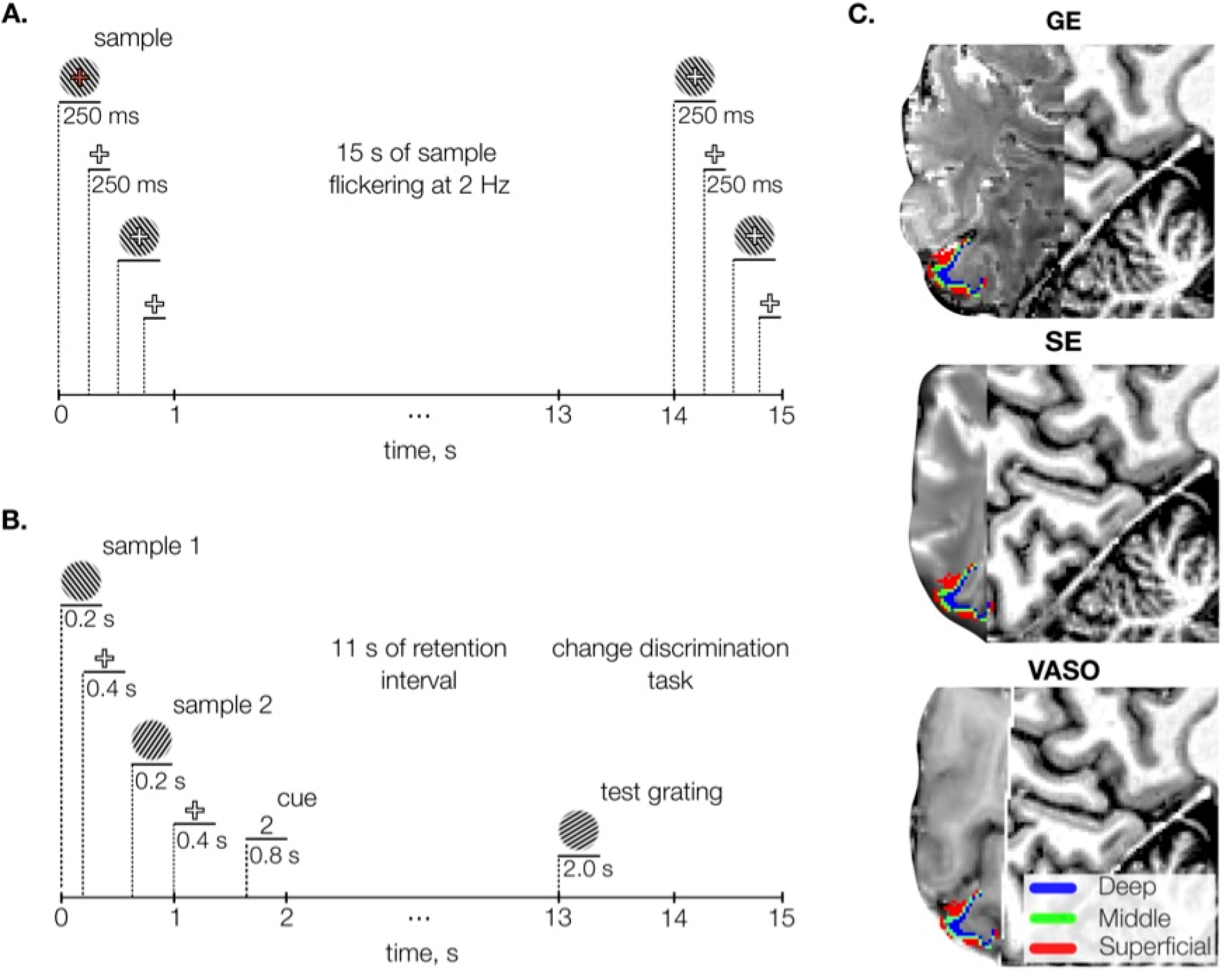
Methods. A. Perceptual task. On each trial participants viewed a sample grating flickering at 2 Hz. They had to press a button when the fixation cross was turning red. **B. Working memory task**. On each trial, participants viewed two sample gratings, then a cue (“1” or “2” in the screen center) indicated which of the grating orientations was to be memorized. After a retention interval of 11 seconds, a probe grating was shown, and participants had 2 seconds to report whether the probe was tilted clockwise or counterclockwise compared to the memorized grating. **C**. Sagittal slice of the T1-weighted anatomical image of a representative participant overlaid with the three average functional images acquired with different MR-sequences (GE-BOLD, SE-BOLD, VASO) and cortical depth bins approximating cortical layers (superficial, middle and deep) from an equi-volumetric model (see Methods). The cortex is mapped within the region of V1 with voxel eccentricity values 1-3°.

#### 2.3.2. Experimental perceptual task

To specifically investigate visual processing dominated by feedforward signals, we included two perceptual runs in the experiment. During these runs, gratings with the two target orientations (25° and 115°) were shown in a pseudo-randomized order. On each trial, one of the grating orientations was shown for 15 seconds, flickering at 2 Hz. The presentation was followed by a fixation cross, which lasted 9 seconds in GE and SE measurements and 10 seconds in VASO measurements. Participants had to monitor the fixation cross for occasional brief changes in color, to which they had to respond with a button press. Overall, in every run we recorded 16 trials evenly split between the two orientations conditions. The perceptual task was performed in the experiment during runs 3 and 6 (out of 9 runs overall). The fixation dot changed 7.4±0.03 times per trial at random time points, leading to approximately 118 changes, to which participants responded on average 75+/-3% (Mean ± SD) of the time. As for the experimental runs, each perceptual task run took 6 minutes 42 seconds for GE and SE measurements and 7 minutes for the VASO measurements.

#### 2.3.3. Experimental working memory task

In the scanner, participants continued to perform the working memory task they were trained on (Figure 1), but with three major changes. First, participants did not receive feedback on their performance. Second, the intertrial interval (ITI) was prolonged to 9 seconds for the measurements with GE and SE sequences and to 10 seconds with VASO sequences. This was done to reduce the temporal overlap between the perceptual trace from the probe grating shown on the previous trial and the current trial. The length of each trial (including ITI) was 24 seconds in the GE and SE sessions and 25 seconds for the VASO sessions. The difference in trial lengths also ensured that every trial was evenly divisible by a fixed number of TRs: with a 3 second TR used for GE and SE measurements, each trial comprised exactly 8 TRs; whereas with 5 seconds TR used for the data acquisition with VASO, each trial comprised exactly 5 TRs. Third, the sample gratings shown at the beginning of each trial were no longer randomly oriented, but always either 25° or 115° orientated away from the vertical axis. We limited the number of possible orientations compared to the training session to increase the signal-to-noise ratio for each grating orientation and thereby enable us to differentiate orientation signals at the level of cortical depth bins.

Every run consisted of 16 trials. In each half of trials, one of the two possible orientations (25° versus 115°) was selected for memorization. Trial order was fully randomized. The experiment consisted of 7 runs, which each lasted 6 minutes 42 seconds for GE and SE and 7 minutes for VASO. The average time for completing the whole experiment in one session was 80-85 minutes.

The anatomical scans for each participant were acquired separately during other studies in MPI Leipzig. None of these scans were older than 6 months.

We invited every participant two times per scanning sequence (GE, SE and VASO) to increase signal-to-noise ratio. Each participant was thus scanned in 6 experimental sessions. The order of the scanning sequences was randomized across the sessions for every subject.

#### 2.3.4. Staircase Procedure

To maintain a sensitive accuracy range across the whole experiment, including the training runs and the fMRI experiment, we used a staircase procedure that adjusted the amount of additional tilt in the probe grating compared to the memorized grating. The initial difference between the memorized orientation and the probe grating was set to 20°. For each correct response in each subsequent trial, the difference between the probe and rotated grating was reduced by 0.5°, making orientation discrimination harder. Conversely, the difference was increased by 2° for each incorrect response, making discrimination easier. We imposed an upper limit of 40° on the orientation difference. Across all the MR-sequences participants’ performance ranged in between (Mean ± SD): 8.8°±5.1°.

### 2.4. Parameters of data acquisition

fMRI was acquired on a 7-T Magnetom (Siemens Healthineers, Germany) whole-body scanner using a single-channel-transmit/32-channel RF receive head coil (Nova Medical Inc, USA). fMRI data were recorded with an isotropic spatial resolution of 0.8 mm using GE-BOLD, SE-BOLD and SS-SI-VASO. For the GE-BOLD protocol we used the CMRR MB sequence (2D EPI readout, repetition time (TR) 3000 ms, echo time (TE) 24 ms, number of slices 50, 78° flip angle, 148 × 148 mm^2^ field of view (FOV), GRAPPA acceleration factor 3, phase partial Fourier 6/8, coronal orientation, F >> H phase encoding direction) (Moeller et al., 2010). For the SE-BOLD, we also used CMRR MB sequence with the following parameters (2D EPI readout, TR 3000 ms, TE 38 ms, number of slices ∼30, 90° flip angle, 148 × 148 mm^2^ FOV, GRAPPA acceleration factor 3, phase partial Fourier 6/8, coronal orientation, F >> H phase encoding direction). The number of slices in SE-BOLD was less than that in GE-BOLD due to specific absorption rates limitations. For the SS-SI-VASO protocol we used the following parameters (3D EPI readout, TR 2837.9 ms, TE 25 ms, TI 650 ms, number of slices 26, 26° flip angle, 133.0 × 133.0 × 20.8 mm^3^ FOV, GRAPPA acceleration factor 3, phase partial Fourier 6/8, orientation T > C-28.2 deg). The number of slices in SS-SI-VASO sequence is less than that in GE-BOLD due to longer TRs and therefore, restricted field of view. Shimming was performed using the standard Siemens procedure. Anatomical data were acquired using a MP2RAGE sequence with 0.7 mm isotropic resolution (3D EPI readout, TR 5000 ms, TE 2.45 ms, TI 900/2750 ms, flip angle 5/3, bandwidth 250 Hz/Px, 224 × 224 × 168 mm^3^ FOV, GRAPPA acceleration factor 2, phase partial Fourier 6/8, base resolution 320, sagittal orientation, scan time 10 min 57 sec).

### 2.5. Functional and anatomical data preprocessing

#### 2.5.1. Bias field correction and segmentation of the anatomical image

The DICOM data were converted to NIfTI format using SPM12 (Statistical Parametric Mapping; Wellcome Center for Human Neuroimaging, University College London). The volumes were bias field-corrected using an SPM-based customized script (Luesebrink, Sciarra, Mattern 2017). To implement cortical depth-specific analysis, we extracted grey matter segmentation for each subject. To do this, first we used the SPM12 segmentation algorithm and then the brainmask was generated by adding up the white matter, grey matter and cerebro-spinal fluid masks. Then we applied the FreeSurfer (version 6.0.0) recon algorithm to perform segmentation of white matter, grey matter, generating their surfaces and a binary brain mask of the cortical ribbon (1 if the voxel falls into the ribbon, 0 otherwise (steps 5-31 of recon-all algorithm)). We ran the recon algorithm on the extracted brainmask from a MP2RAGE image with a ‘-hires’ flag for the data with resolution higher than 1 mm (Zaretskaya et al., 2017). After running the recon algorithm, the Freesurfer-generated grey and white matter segmentations were visually inspected in each participant, the borders between CSF and grey matter as well as grey matter and white matter were manually corrected within the region corresponding to the field of view of functional scans.

#### 2.5.2. Cortical depth and ROI definition

The grey matter segmentation acquired with Freesurfer was further utilized to obtain cortical depth-specific bins. Deep, middle and superficial bins were constructed using an equi-volumetric model (Waehnert et al., 2014; Huntenburg et al., 2018). In order to analyze depth-specific activity in early visual areas, we applied a probabilistic surface-based anatomical atlas (Benson et al., 2014) to reconstruct the surfaces of area V1 for each subject. This is an atlas of the visual field representation (eccentricity and polar angle), and eccentricity values were used to select the foveal sub-part of the surface excluding the area occupied with the fixation cross (1-3°). The extracted surface ROI (V1) was then projected into the volume space and intersected with the predefined cortical bins. In this way, we obtained the V1 ROI in the Freesurfer anatomical space at three predefined cortical depths.

#### 2.5.3. Functional data preprocessing

Functional volumes were realigned to the first volume of the middle run using SPM software. In the data acquired with VASO sequence, the nulled and not nulled frames were realigned separately (Finn et al., 2019), the not-nulled volumes were interpolated using 7th order spline function to correspond to the acquisition time of the nulled frames (the nulled volumes did not undergo temporal interpolation in our study, in order to reduce the sharing of informational content across time points). Motion traces of nulled and not-nulled frames were then visually inspected to ensure good overlap of these contrasts. The nulled data was then corrected with the not-nulled volumes using the dynamic division method (Finn et al., 2019).

After the realignment, we ensured that the functional runs acquired with each scanning sequence were well aligned with each other in each participant. This is required for multivariate pattern analyses of high-resolution fMRI data. For this we computed inter-run spatial cross-correlations of the signal intensities of the functional volumes. The resulting average spatial correlation of experimental runs was very high: (Mean ± SD) 0.97±0.03 in GE-BOLD sessions, 0.986±0.01 in SE-BOLD sessions and 0.99±0.004 in SS-SI-VASO sessions. Further, functional-anatomical alignments were checked visually to ensure that the functional scans were well aligned to the anatomical image at the location of the ROIs.

After that, the data acquired with all the three sequences was high-pass filtered (removing signal with f<1/128 Hz). Before classification analysis, the functional time series of every voxel within the ROI was z scored to correct for the scaling differences in voxel intensities within every run (Lawrence et al., 2018).

#### 2.5.4. Registration of fMRI and anatomical data

We linearly coregistered the extracted ROIs with predefined cortical depth bins to the EPI volumes within each subject using the Symmetric Normalization (SyN) algorithm of ANTs (Avants et al., 2008). Specifically, first, the T1-weighted anatomical image was registered using linear interpolation to the EPI volume averaged over all the functional runs. Next, we registered the ROIs with the predefined cortical depths to the EPI volume using nearest neighbor interpolation and by applying the coordinate mapping (with the voxel size resampled to the functional runs (0.8 isotropic)) obtained in the previous step. The resulting ROIs included the following number of voxels per cortical depth and MR-sequence (Mean ± SD): GE-BOLD (M_deep_=1170.8±90, M_mid_=1083±62, M_super_=1170.2±109), SE-BOLD (M_deep_=1026.1±197, M_mid_=955.1±158, M_super_=1035±216), SS-SI-VASO (M_deep_=1120.8±97, M_mid_=1037±95, M_super_=1120±143).

### 2.6. Multivariate pattern analysis

#### 2.6.1. Data extraction

Multivariate pattern analysis (MVPA) was performed in each subject and experimental session separately. To prepare the EPI data for the MVPA, we first extracted activity patterns for a V1 ROI with the predefined cortical depths from the functional images in the experimental task runs. Specifically, in each run, we extracted voxel-wise activation values for the 2 oriented grating conditions (25° and 115°) and 8 trials for each condition across trial TRs starting at trial onset.

#### 2.6.2. Classification

MVPA was carried out using linear support vector machines (SVMs; libsvm: http://www.csie.ntu.edu. tw/∼cjlin/libsvm/) with a fixed cost parameter (c=1). We performed classification both across all the voxels in the full grey matter ribbon of V1 (Supplementary Figure 1) as well as separately at each cortical depth of V1. To this end, we trained the SVM classifiers on multi-voxel response patterns from all the experimental runs within each experimental session, leaving out one run (i.e., using leave-one-out cross validation), to discriminate between the 2 oriented gratings for each time point in the trial. All trials in the training set were utilized for the classifier training (8 trials per orientation and training set in perception task and 48 trials per orientation and training set in working memory runs). Next, we tested the SVM classifier using the trials of the left-out run (8 trials in both tasks). The classifier was trained on each time point in the trial using the data from the training set and tested on the corresponding time point in the test set. As a result, we extracted decoding accuracy for every TR of all the runs in the main experiment (chance level 50%). The results were organized in the form of the matrix for further statistical testing: participants (4) x experimental runs (number of runs depended on the working memory (7) or perception task (2)) x TRs (number of TRs depended on the scanning sequence (GE - 8; SE - 8; VASO - 5)).

#### 2.6.3. Statistical testing

For the statistical assessment of feedforward and feedback effects, we preselected the time interval in the trial of a respective task. We estimated the feedforward effect by averaging over the classification accuracy obtained during the 12 seconds interval following stimulus presentation in the perception task trials. In GE-and SE-acquired data, the measurements at 6, 9 and 12 seconds were included in this analysis (Supplementary Figure 1). The first time point in the trial (after 3 seconds) was excluded as uninformative, as it was too close to the stimulus presentation to carry reliable orientation information. In the VASO data, the classification accuracies obtained at 5 and 10 seconds in the trial were averaged and underwent further statistical testing. Here, we did not exclude the first time point (after 5s), since this later time point was likely to already contain information about the stimulus presentation.

For the statistical assessment of feedback contents, we preselected the critical time interval based on previous studies (Harrison & Tong, 2009; Albers et al., 2013; Rademaker, Chunkaras & Serences, 2019) where working memory representations could be decoded starting from 6 seconds after stimulus onset. For GE and SE measurements, the critical interval in our study included measurements at 6, 9, 12 and 15 seconds, that is, all the time points starting from 6 seconds until one time point after the probe grating onset (at 13 seconds) (Supplementary Figure 1). We also included the time point after the probe grating presentation, since this measurement was too close to the presentation of the probe grating to be contaminated by it (Iamshchinina et al., 2021). For the statistical analysis of VASO-based estimates, we included the time points starting from 6 seconds or later, which resulted in measurements at 10 and 15 seconds.

To test whether decoding of orientation in feedforward and feedback signals was significantly above chance and to compare the decoding between cortical depth bins, we used a linear mixed modeling approach (fitlme function, Matlab, The Mathworks Inc, 2014). First, in order to obtain a baseline against which the presence of the signal could be tested, we generated permuted null distributions of classification accuracies for each participant, timepoint and sequence. On each permutation, we reshuffled orientation labels for all trials before performing the classification (1,000 iterations with shuffled data labels). Then, we averaged the classification results of all the permutations and compared them to the original classification results (the empirical effect). The feedforward and feedback conditions were estimated separately due to substantial differences in the paradigms and trial time intervals chosen. The data from all the runs of all the participants were concatenated within each experimental session without averaging (i.e., 16 data points in the perception task (4 subjects X 2 sessions per imaging method X 2 experimental runs) and 56 data points in the working memory task (4 subjects X 2 sessions per imaging method X 7 experimental runs)). The linear mixed effects model included an intercept, classification accuracy as a response variable, cortical depth variable as a fixed effects portion of the model (categorical predictor), participants number as a grouping variable with the random-effects terms for intercept and cortical depth specified separately (assuming no correlation between them): classification accuracy ∼1+depth_effect + (1|participants) + (depth_effect-1|participants).

## 3 Results

In this study, we compared the performance of three MR-sequences (GE, SE and VASO) in distinguishing feedforward and feedback signals in cortical depth of area V1. Using each MR-sequence and 7T fMRI, we measured feedforward and feedback-dominated information emerging during perception and memorization of orientated gratings, respectively (Figure 1). We collected data from 4 participants, where each participant undertook two sessions per scanning method. Signal estimation was performed in separate grey matter bins (superficial, middle and deep) extracted with an equi-volumetric model (see Methods). For each depth bin, we trained support vector machine classifiers to differentiate multi-voxel response patterns evoked by the different grating orientations. For this analysis, we focused on time points which were expected to carry robust perceptual signals (measurements at 5 and 10 seconds in VASO measurements and for other sequences: 6, 9 and 12 seconds) or memory signals (measurements at 10 and 15 seconds in VASO measurements and for other sequences: 6, 9, 12 and 15 seconds). By comparing classification in these time windows between the grey matter bins, we establish which of the MR-sequences uncovers differences in feedforward and feedback signals across cortical depth.

### 3.1 Feedforward signals across cortical depth in V1

We examined the cortical profile of orientation-selective activity measured during the perception task with the three MR-sequences. Firstly, we established whether the different perceived orientations could be successfully decoded at different cortical depth. Secondly, following the predictions based on previous animal and human work (Rockland & Pandya, 1979; Lawrence et al., 2019), we hypothesized that the middle cortical bin represents the perceived contents more strongly than the outer cortical bins averaged together.

Analysis of the GE-BOLD data yielded above-chance classification results in the middle cortical bin (t(26)=2.6, p=0.01) but not in the other bins (p>0.1). There was also no significant difference between the cortical bins (p>0.4) (Figure 2A).

**Figure 2.**
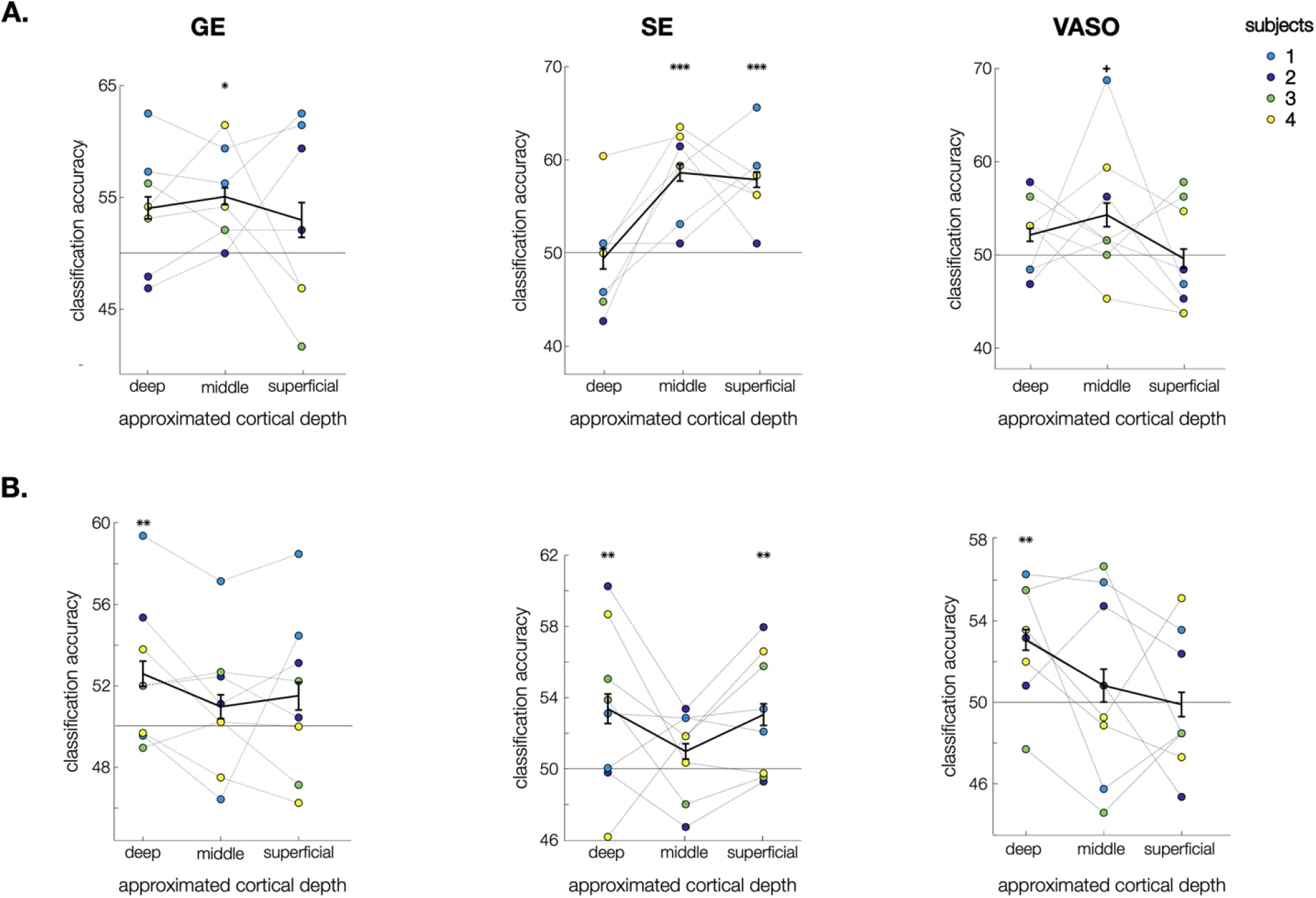
Results. **A**. Classification accuracy for decoding of perceived grating orientation in V1 measured with GE-BOLD, SE-BOLD and VASO. Above-chance classification results for perceived orientation were found in the middle cortical bin in all the sequences and at superficial depth for SE-BOLD. Significant differentiation across cortical depth only emerged for SE-BOLD, where decoding was stronger in the middle bin than in the other two bin combined. **B**. Classification accuracy for decoding of memorized grating orientation. Above-chance classification results for memorized orientation were found in the deep cortical bin in all the sequences and in the deep and superficial cortical bins for SE-BOLD. A significant differentiation between the middle and outer depth bins was again only achieved using SE-BOLD. The colored dots represent experimental sessions (2) for each participant (4). The data points are to show the correspondence between two separate sessions within each participant. Note that the statistical analysis was performed using experimental runs unaveraged within each of the sessions. All error bars denote standard error of mean within participants. Represented p-values are fixed effects coefficients estimated with a t-test for each cortical depth (uncorrected) +: p<0.07, *: p<0.05, **: p < 0.01, ***: p < 0.001.

Analysis of the SE-BOLD data revealed a highly significant above-chance classification results in the middle (t(26)=5.0, p<0.001) and superficial bins (t(26)=4.2, p<0.001) but not in the deep bin (p=0.7). In accordance with our hypothesis about a differential representation of the feedforward signal, we observed a higher classification accuracy for perceived orientation in the middle bin than in the superficial and deep bins averaged together (t(26)=2.7, p=0.01). Unpacking this effect, we found a higher classification accuracy for perceived orientation in the middle bin than in the deep bin (t(26)=4.9, p<0.001), with no difference between the middle and the superficial bins (p=0.7).

Finally, analysis of the VASO data yielded above-chance classification results for perceived orientation in the middle cortical bin (on the trend-level; t(30)=2.0 p=0.06) and not in the other bins (p>0.2), with no significant differences between the cortical bins (p>0.3).

Overall, we localized feedforward signals at the middle cortical depth bin of V1 for all the MR-sequences. However, a more strongly represented feedforward effect in the middle cortical bin compared to the outer cortical bins was only observed in the data acquired with SE-BOLD, but not with the other sequences. This differential feedforward representation for SE-BOLD was mainly driven by a difference between the middle and deep cortical bins.

### 3.2 Feedback signals across cortical depth in V1

Next, we examined differences in revealing feedback representations in human V1 across the three scanning sequences. We analyzed the orientation-selective activity acquired during the working memory task focusing our analysis on the retention time interval (see Methods). As for the analysis of the perception signal, we aimed to establish which cortical bins carry the representation of the memorized item. Firstly, we established whether the different memorized orientations could be successfully decoded at different cortical depth. Secondly, following our predictions about the layer-specific distribution of feedback signals, we tested if any of the outer cortical bins represented the memorized contents more strongly than the middle cortical bin (Figure 2B).

Analysis of the GE-BOLD data revealed above-chance classification results in the deep cortical bin (t(104)=2.8, p=0.007; Figure 2B) but not in the other bins (p>0.2), with no significant difference between the bins (p>0.3).

Analysis of the SE-BOLD data yielded above-chance classification in both the deep (t(102)=2.7, p=0.008) and superficial (t(102)=2.9, p=0.004) bins, but not in the middle bin (p>0.26). Comparing feedback representations across cortical depth revealed that classification accuracy for memorized orientation in the outer bins averaged together was higher than the in middle bin (at the trend level; t(102)=1.8, p=0.07). Pairwise comparison between all three cortical bins showed higher classification accuracy for memorized orientation in the superficial bin than in the middle bin at the level of trend (t(102)=1.9, p<0.06) with no difference between the rest of the bins (p>0.15).

Finally, analysis of the VASO data revealed above-chance classification results for memorized orientation only in the deep cortical bin (t(110) = 2.6, p = 0.009) but not in the other cortical bins (p>0.5), with no difference between the cortical bins (p>0.6).

Overall, we observed feedback signals in the deep cortical bin of V1 for all the MR-sequences. However, we only observed a differential representation of feedback signals across the middle and outer cortical bins of V1 when the data was acquired with SE-BOLD, but not with the other sequences. This differential feedback representation for SE-BOLD was mainly driven by a difference between the superficial and middle cortical bins.

## 4. Discussion

In the present study, we investigated depth-dependent separation of feedforward and feedback signals at 7T fMRI using three MR-sequences: GE-BOLD, SE-BOLD and SS-SI-VASO. We used multivariate pattern classification to read out information about the contents of feedforward and feedback information across cortical depth in area V1.

For all three sequences, we were able to decode feedforward signals from the middle depth bin of area V1, while feedback could be read out from the deep cortical bin. This consistency indicates that the widely used GE-BOLD (Muckli et al., 2015; Kok et al., 2016;

Lawrence et al., 2018; Bergmann et al., 2019; Aitkens et al., 2021; Iamshchinina et al., 2021) is a viable method for decoding information from cortical depth bins. This suggests that despite its lower spatial specificity than SE-BOLD or VASO (Yacoub et al., 2003; Duong et al., 2003; Olman et al., 2010; Huber et al., 2014), the more sensitive GE-BOLD measurements can detect depth-specific signals in V1.

From the three sequences SE-BOLD stood out by being the only sequence in our study that yielded a statistically reliable depth-specific separation of the feedforward and feedback signals, where feedforward representation was significantly stronger in the middle bin compared to the outer bins and feedback representation was stronger in the outer bins compared to the middle bin. These results align well with previous animal research and 7T work in humans (Kok et al., 2016; Aitken et al., 2021; Bergmann et al., 2019) and highlight SE-BOLD as a particularly interesting MR-sequence for establishing layer-specific effects in experiments of the kind used here, through its favorable trade-off between sensitivity and spatial specificity.

The depth differentiation was mainly reflected in differences between the middle and deep bins. The estimates obtained in the superficial depth bin were found to be less consistent across the MR-sequences. This could be related to a bias towards the surface veins observed in the SE-BOLD measurements (Uludag, Mueller-Bierl, Ugurbil, 2009) which could have impacted the effects obtained at the superficial cortical depth. Alternatively, however, the generally lower classification obtained with VASO may have missed signals at the superficial depth. Future studies need to establish whether feedback signals at the superficial cortical depth can be disentangled from nonspecific contributions such as surface-vein effects. For this, they could use model-based approaches (Markuerkiaga, Bath & Norris, 2016; Havlicek & Uludag, 2020; Markuardt et al., 2017; Huang et al., 2021) or combine high-field MRI with different imaging methods (Kashyap et al., 2020; Huber et al., 2018).

While SE-BOLD yielded the clearest depth separation in the current benchmark study, several other aspects need to be taken into consideration when deciding between imaging protocols for future studies. First, SE-BOLD sequences typically yield reduced BOLD sensitivity compared to GE-BOLD (Boyacioglu et al., 2014). In this context, it is worth noting that our study employed a design with few participants with long acquisitions each, which may be the experimental regime particularly suited for SE-BOLD. Second, they are affected by unspecific factors, such as residual venous signal contributions, sensitivity to B1-inhomogeneity, which can dramatically affect the quality of acquired data, and are limited by specific absorption rates (SAR), which can drastically reduce the number of slices that can be acquired (van der Zwaag et al., 2014; Marques & Norris, 2018). Given these limitations, future studies could employ improved protocols for SE-BOLD looking to overcome currently present challenges (Norris et al., 2011; Mugler, 2014; Gagoski et al., 2015; Han et al., 2021).

In our experiment VASO data showed low decoding accuracy for feedforward and feedback signals (Supplementary Figure 2 and 3). This unfavorable outcome could be partially due to lower temporal resolution of VASO measurements and thus lower statistical power in its analysis compared to other MR-sequences. Specifically in our study, the selection of time intervals for the assessment of feedforward and feedback cortical profiles was more challenging for VASO, because each TR covered larger temporal chunks within the trial (see Supplementary Figure 3).

In sum, whether GE-BOLD, SE-BOLD or VASO are the most appropriate choice for dissociating feedback from feedforward information will in practice depend on multiple factors of the experimental context. Our study contributes to mapping out the choice space by providing a single point in this space through benchmarking for a typical cognitive neuroscience experiment – however, future investigation is needed to populate this choice space further, with different protocols and paradigms.

## 5. Data/code availability statement

The data and the code will be available upon acceptance of the manuscript for publica-tion.

## 7. Acknowledgments

We thank the University of Minnesota Center for Magnetic Resonance Research for the provision of the multiband EPI sequence software. PI is supported by the Berlin School of Mind and Brain PhD scholarship. DK and RMC are supported by Deutsche Forschungsgemeinschaft (DFG) grants (KA4683/2-1, CI241/1-1, CI241/3-1, CI241/7-1). RMC is supported by a European Research Council Starting Grant (ERC-2018-StG). NW received funding from the European Research Council under the European Union’s Seventh Framework Programme (FP7/2007-2013) / ERC grant agreement n° 616905; from the European Union’s Horizon 2020 research and innovation programme under the grant agreement No 681094; from the BMBF (01EW1711A & B) in the framework of ERA-NET NEURON. Computing resources were provided by the high-performance computing facilities at ZEDAT, Freie Universität Berlin.

## 8. Author contributions

P.I., D.K. and R.M.C. designed the study, D.K., R.M.C. and N.W. supervised the study, P.I., D.H., R.T. acquired data. P.I., D.K. and D.H. analyzed the data. P.I., D.K., R.M.C. wrote the original draft of the manuscript, R.T., D.H., N.W. reviewed and edited the manuscript.

---

**Supplementary Figure 1.**
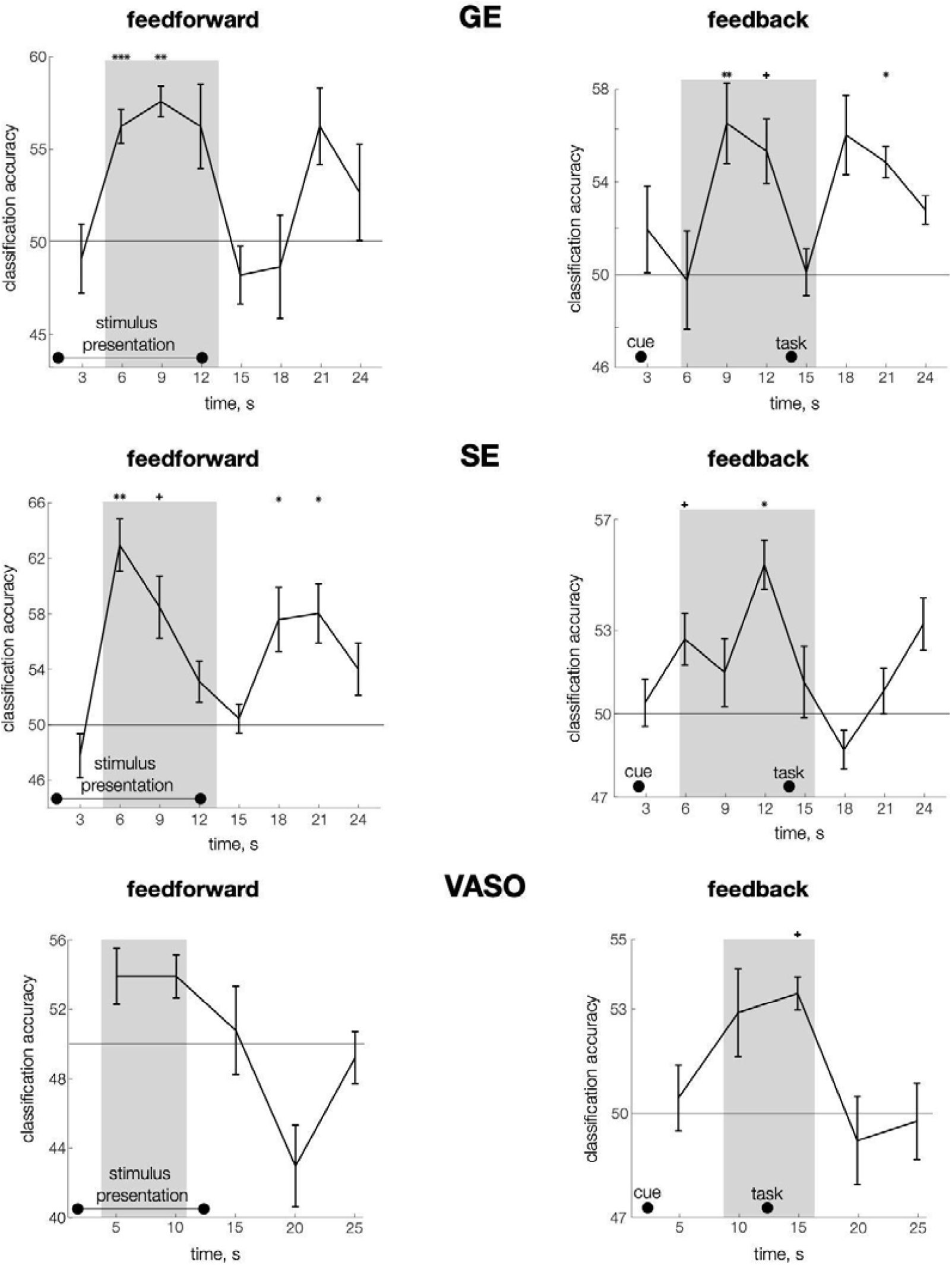
**Left column**.Classification time series of feedforward (perceived) contents (i.e. grating orientations) extracted from the full grey matter ribbon (all voxels included in classification analysis) and represented for every MR sequence. During the stimulus presentation phase of the trial, a significant representation of perceived grating orientation emerged in the analysis of GE-and SE-acquired data (t(26)=2.96, p=0.007, t(26)=3.1, p=0.004 for GE and SE, respectively). VASO data yielded above-chance classification results for perceived orientation at the level of trend (t(30)=1.9, p=0.07). Thus, we established the presence of the feedforward signal in the data acquired with all the sequences. Somewhat weaker representational strength obtained from VASO measurements could be due to coarser time resolution and consequently the inclusion of less time points in this analysis. **Right column**. Same as the left column but for the feedback (working memory) classification results. The analysis based on all the voxels in the V1 ROI showed an above-chance classification results for memorized orientation in all the sequences (t(104)=1.9, p=0.06, t(102)=2.8, p=0.005, t(110)=2.6, p=0.01 for GE, SE and VASO sequences, respectively). Thus, our results indicate that it is possible to decode working memory representations from the data acquired not only with GE 7T fMRI (Lawrence et al., 2018) but also SE-BOLD and VASO scanning methods.

**Supplementary Figure 2.**
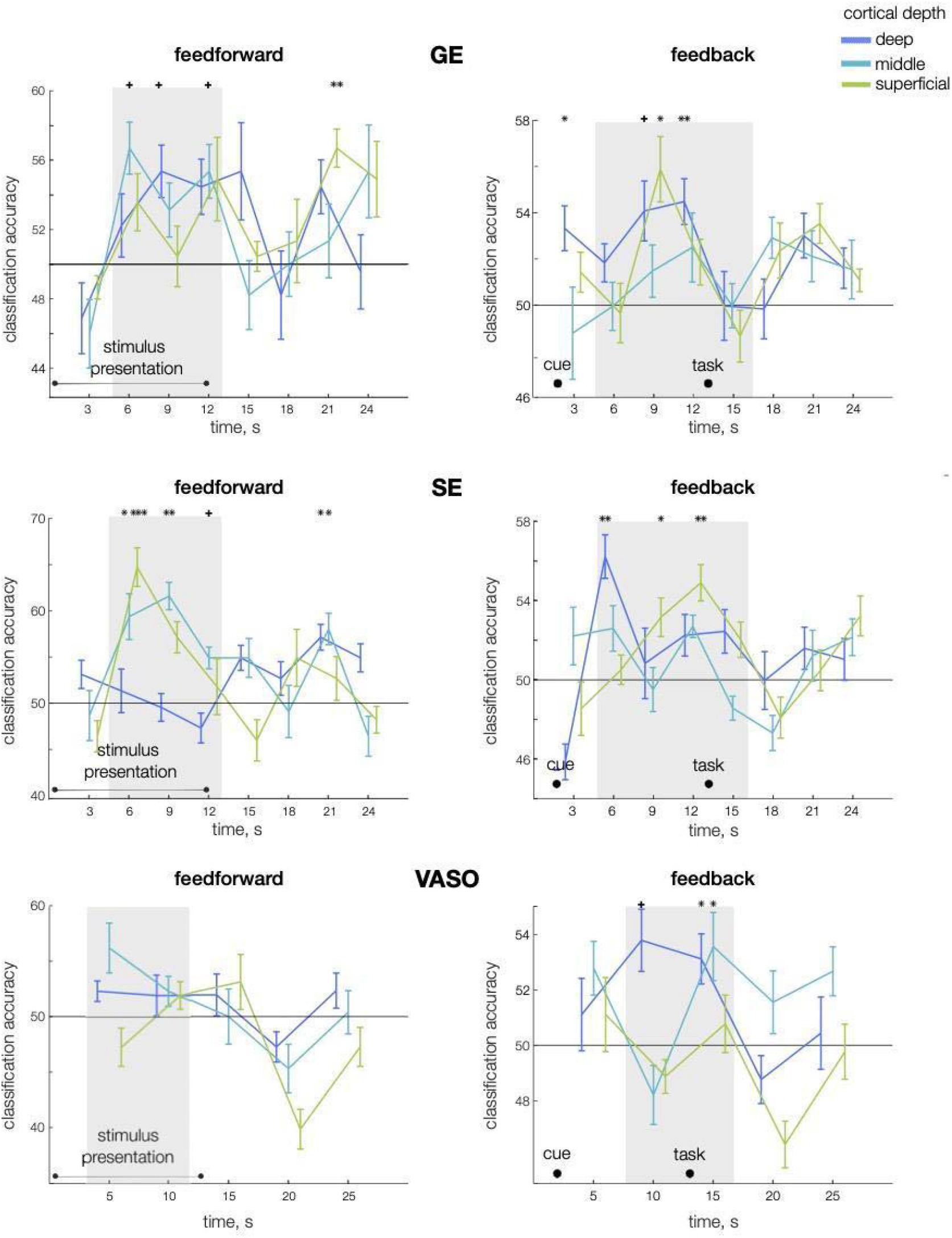
**Left column.**Time series of classification accuracy for feedforward contents (perceived orientation) extracted from the three approximated cortical depth bins and represented for every MR sequence. **Right column**. Same as the left column but for the feedback contents (memorized orientation).

